# Crystal Structures of Inhibitor-Bound Main Protease from Delta- and Gamma-Coronaviruses

**DOI:** 10.1101/2023.02.01.526623

**Authors:** Sarah N. Zvornicanin, Ala M. Shaqra, Qiu Yu J. Huang, Elizabeth Ornelas, Mallika Moghe, Mark Knapp, Stephanie Moquin, Dustin Dovala, Celia A. Schiffer, Nese Kurt Yilmaz

## Abstract

With the spread of SARS-CoV-2 throughout the globe to cause the COVID-19 pandemic, the threat of zoonotic transmissions of coronaviruses (CoV) has become even more evident. As human infections have been caused by alpha- and beta-CoVs, structural characterization and inhibitor design mostly focused on these two genera. However, viruses from the delta and gamma genera also infect mammals and pose potential zoonotic transmission threat. Here, we determined the inhibitor-bound crystal structures of the main protease (M^pro^) from the delta-CoV porcine HKU15 and gamma-CoV SW1 from beluga whale. Comparison with the apo structure of SW1 M^pro^, which we also present here, enabled identifying structural arrangements upon inhibitor binding at the active site. The binding modes and interactions of two covalent inhibitors, PF-00835231 (lufotrelvir) bound to HKU15 and GC376 bound to SW1 M^pro^, reveal features that may be leveraged to target diverse coronaviruses and toward structure-based design of pan-CoV inhibitors.

## Introduction

Viral species from the coronavirus (CoV) family can cause severe respiratory, enteric and hepatic disease, both in animals and humans. Wildlife, especially bats, harbor a diverse pool of coronavirus species that cross-over to other animals including mammals [1]. Further evolution of the virus to adapt to mammalian host increases the likelihood of zoonotic transmission to humans. Three such transmissions caused major outbreaks in humans, namely SARS-CoV-1, MERS, and most recently SARS-CoV-2, which are thought to have originated in bats but transmitted through civets, camels, and possibly pangolins as intermediate hosts to humans [2-4]. Thus, investigating coronaviruses from various hosts, especially other mammals, is important to evaluate the natural diversity and evolution of these viruses toward readiness for future cross-species transmissions and potential outbreaks.

Despite the diversity and rapid evolution of coronavirus species, as exemplified in SARS-CoV-2 during the COVID-19 pandemic, certain elements of the viral genome are highly conserved. While the spike protein shows extreme diversity and quick adaptation to thwart antibody responses and vaccine effectiveness, the 3C-like main protease (M^pro^) has remained relatively unchanged. In fact, M^pro^ from SARS-CoV-1 and -2 are 96% identical. SARS-CoV-2 M^pro^ also shares the same overall fold and active site with 3C and 3C-like proteases from various coronavirus species and even other viral families [5,6]. This similarity prompted the exploration of potential pan-3C inhibitors that leverage the conserved substrate specificity at the P1 and P2 positions, to target the conserved features of the active site. These similarities have enabled repurposing 3C protease and M^pro^ inhibitors to inhibit SARS-CoV-2 and the unprecedented rapid clinical development of COVID-19 antivirals [7,8]. One such broad-spectrum inhibitor is GC376, which inhibits many picornavirus-like species including rhinoviruses, enteroviruses, and coronaviruses [9]. GC376 is a bisulfite derivative prodrug that converts to the active aldehyde form GC373 and covalently attaches to the catalytic Cys of the protease. This dipeptidyl compound was first reported to inhibit feline alpha-CoV FIPV, and its cocrystal structure was solved with M^pro^ of FIPV as well as the porcine alpha-CoV TGEV [9,10] (PDB IDs: 4F49, 7SMV). GC376 also inhibits the M^pro^ of beta-CoVs, including SARS-CoV-2, with nanomolar Ki values. The co-crystal structure for this complex was determined by multiple groups during the COVID-19 pandemic [11-13] (PDB IDs: 6WTJ, 6WTT, 7JSU), and M^pro^ inhibition by GC376 was further characterized to be covalent but reversible [14]. Another repurposed compound that progressed into clinical trials as lufotrelvir is the hydroxymethylketone-based covalent inhibitor PF-00835231 [7]. This phosphate prodrug converts to its active form PF-07304814 and attaches irreversibly to the active Cys (PDB ID: 6XHM). Both these inhibitors were structurally characterized against M^pro^ of alpha-and beta-CoVs, but their binding mode to more divergent delta- and gamma-CoVs M^pro^ variants has not been described.

Viral species from the delta genus of coronaviruses have mostly been isolated from birds, but they can also infect mammals as exemplified by the porcine HKU15. The gamma-CoVs were more recently identified as a genus, with the discovery of a novel and highly divergent coronavirus (named SW1) in the liver of a captive beluga whale [15]. Thus delta- and gamma-CoVs have been shown to infect mammals that live in proximity to humans. The M^pro^ amino acid sequence of HKU15 and SW1 are less than 50% identical to those from beta-CoVs as well as to each other (Figure S1). The crystal structure of HKU15 M^pro^ has recently been determined with a relatively large peptidomimetic inhibitor [16] but how potential pan-CoV inhibitors bind to HKU15 is not known. Additionally, apart from the avian infectious bronchitis virus (IBV) [17], structural characterization of M^pro^ from gamma-CoVs is severely lacking.

In this work, we report the crystal structures of M^pro^ from the gamma-CoV SW1 in both apo and inhibitor-bound form, as well as the crystal structure of the delta-CoV HKU15 M^pro^ in complex with PF-00835231. These structures enabled characterization and comparison of the main viral protease from these divergent mammal-infecting coronaviruses and identification of active site features that might be critical for the design of pan-coronaviral inhibitors for future potential outbreaks.

## Materials and Methods

### Protein expression and purification of M^pro^

M^pro^ sequences were cloned into the pETite expression plasmid using standard techniques. The constructs included an N-terminal polyhistidine – SUMO tag which was used for affinity purification and to prevent the toxic activity of the protein from impeding bacterial growth.

HI-Control™ BL21(DE3) (Lucigen) cells were transformed with plasmid and single colonies were picked to start overnight cultures in LB medium supplemented with 50 μg/mL kanamycin at 37 °C. These cultures were used to inoculate 1 L cultures in TB medium containing 50 mM sodium phosphate (pH 7.0) and kanamycin (50 μg/mL). When the OD600 value reached ∼1.5–2.0, 0.5 mM IPTG was added to induce protein expression and the cell culture was further incubated overnight at 19 °C. Cells were harvested by centrifugation at 5,000 rpm for 30 min, resuspended in Lysis Buffer (50 mM Tris-HCl (pH 8.0), 400 mM NaCl, 1 mM TCEP) and lysed by a cell disruptor. The lysate was clarified by centrifugation at 45,000 x g for 30 min. The supernatant was then subjected to IMAC purification by loading onto a HisTrap FF column (GE Healthcare) equilibrated with lysis buffer, washed with lysis buffer until the A280 stabilized, and followed by elution using a linear gradient of Lysis Buffer supplemented with 0–500 mM imidazole over 40 column volumes. The fractions containing M^pro^ were pooled and treated with ULP1 to remove the His-SUMO tag and produce an authentic

N-terminus. Cleavage occurred overnight at room temperature while dialyzing into 3 L of Lysis Buffer (0 mM imidazole). We found prolonged exposure to 4 °C often led to substantial precipitation of the protein, which is why dialysis occurred at room temperature. Full cleavage was confirmed by ESI-LC/MS and the cleaved protein was again subjected to IMAC purification by flowing through 5 mL of Ni-NTA resin pre-equilibrated with Lysis Buffer. The supernatant was collected and concentrated to approximately 5 mL before further pufication by size-exclusion chromatography (SEC). SEC purification was performed on a HiLoad 16/60 Superdex 75 column (GE Healthcare) pre-equilibrated with 25 mM HEPES pH 7.5, 150 mM NaCl, and 1 mM TCEP. Pure fractions were pooled, concentrated to 10–15 mg/mL, aliquoted, and stored at –70 °C. The final yield from 1 L of bacteria expressing M^pro^ varied between 40 and 70 mg.

### Protein Crystallization

For co-crystallization, protease–small molecule complexes were assembled by incubating 6 mg/mL of each M^pro^ with 10-fold molar excess of ligand for 1 hour at room temperature. Large crystals were obtained with 10–20 % (w/v) PEG 3350, 0.2 M NaCl, and 0.1 M Bis-Tris Methane pH 5.5 by hanging drop vapor diffusion in pre-greased VDX trays (Hampton Research) at room temperature. Varying the protein to mother liquor ratios (1 mL:2 mL, 2 mL:2mL, 3 mL:2mL) per well helped obtain large, diffraction quality crystals. Cocrystals of M^pro^ with PF-00835231 appeared overnight and grew large enough for in-house data collection within 1 week. Crystals of SW1 and HKU15 M^pro^ complexes required 2–4 weeks of growth time to reach a sufficient size for data collection.

For apo crystals, SW1 M^pro^ was screened against multiple crystallization conditions at 14.3 mg/mL (25 mM HEPES pH 7.5, 150 mM NaCl, 1 mM TCEP). SW1 M^pro^ protein was supplemented with 1 mM EDTA and incubated for 10 min before drop setup. Best crystals obtained grew at 18°C from JCSG+ Screen (Nextal Biotechnologies) composed of 25% Peg3350, 0.2 M Ammonium Sulfate and 0.1 M Bis-Tris pH 5.5. The crystals reproduced easily and were grown by hanging drop vapor diffusion, 1:1 mL volume ratio. After 24 h, crystals were harvested and cryo-protected using 20% glycerol and well solution, flash-frozen in liquid nitrogen for data collection. Diffraction data was collected for apo SW1 M^pro^ crystals at the Advance Light Source beamline 5.0.2.

### Data Collection and Structure Determination

Data collection for co-crystals was performed using a MicroMax-007HF x-ray generator equipped with a HyPix-6000HE detector (Rigaku Corporation) at the University of Massachusetts Chan Medical School, Crystallography and Structure Based Drug Design Core Facility. As data was collected under cryogenic conditions (100 K), crystals were protected from freeze damage by a quick soak in crystallization solution supplemented with 25 % glycerol. Diffraction data were indexed, integrated, and scaled using CrysALISPROPX (Rigaku Corporation). Prior to analysis, data quality assessment was done using Xtriage [18]. The structures were solved by molecular replacement using PHASER [19] with 7L0D [20] for the SARS-CoV-2 M^pro^–PF-00835231 structure, SW1 apo M^pro^ (8FWX) to solve the SW1 M^pro^–GC376 structure, and 7WKU [16] to solve the HKU15 M^pro^–PF-00835231 structure. In each case, models for molecular replacement were prepared by removing ligands, structural and bulk solvent waters, and cryoprotectant molecules. SDF files of PF-00835231 (PDB: V2M), N-[(2S)-1-({(2S,3S)-3,4-dihydroxy-1-[(3S)-2-oxopyrrolidin-3-yl]butan-2-yl}amino)-4-methyl-1-oxopentan-2-yl]-4-methoxy-1H-indole-2-carboxamide, and GC376 (PDB: UED), N∼2∼-[(benzyloxy)carbonyl]-N-{(2S)-1-hydroxy-3-[(3S)-2-oxopyrrolidin-3-yl]propan-2-yl}-L-leucinamide were obtained from the Protein Data Bank. These models reflect the ligands in the covalently bound state. Using V2M and UED SDF files as inputs, eLBOW [21] generated ligand atomic position and constraint cif files and corresponding PDB coordinates necessary to fit the small molecules into their respective densities. COOT [22] was used for modeling building while iterative rounds of refinement were performed using PHENIX [18]. Due to merohedral twinning, the HKU15 M^pro^–PF-00835231 structure was refined using twin law -h-k, k, -l. To limit bias throughout the refinement process, five percent of the data was reserved for the free R-value calculation [23]. Model quality assurance was assessed with MolProbity [24] prior to PDB deposition [25,26].

Structure analysis, superposition and figure generation was done using PyMOL [27]. Hydrogen bonds and van der Waals contacts were determined as described previously [28]. X-ray data collection and crystallographic refinement statistics are presented in the Supporting Information (Table S1).

## Results

The crystal structures of M^pro^ from gamma-CoV SW1 (from beluga whale) were determined with a bound covalent inhibitor at the active site, and in the apo form with no inhibitor. Both structures were solved in the P21 space group with the apo form at 2.57 and the inhibitor-bound structure at sub-2 Å resolution. We also determined the inhibitor-bound structure of M^pro^ to 2.45 Å resolution from the porcine delta-CoV HKU15, and the same inhibitor (PF-00835231) bound to M^pro^ of SARS-CoV-2 for direct comparison. All 4 structures were solved with the dimer in the asymmetric unit. The cocrystal structures had full occupancy of inhibitor at both active sites, covalently bound to the catalytic cysteine. Crystallographic data collection and refinement statistics are presented in Table S1.

### Crystal structure of inhibitor-bound gamma-coronavirus SW1 M^pro^

The crystal structure of M^pro^ from beluga whale gamma-CoV SW1 shares the same overall fold of previously reported coronavirus M^pro^ structures, with the C-terminal helical Domain III facilitating dimerization, and the active site between Domains I and II in both protomers accessible for ligand binding (Figure 1). The bisulfite prodrug GC376 converted to the aldehyde form (GC373) and attached covalently to the catalytic Cys142; both active sites were fully occupied by the covalent inhibitor. As previously seen in other M^pro^ cocrystal structures [9,29], the attachment of the aldehyde warhead can result in either (S) or (R)-configuration of the hemithioacetal to orient the hydroxyl group. The electron density was best fit with a 50/50 occupancy of (S)- and (R) configurations at both active sites (in the figures, only the (S) configuration is displayed for clarity).

**Figure 1.**
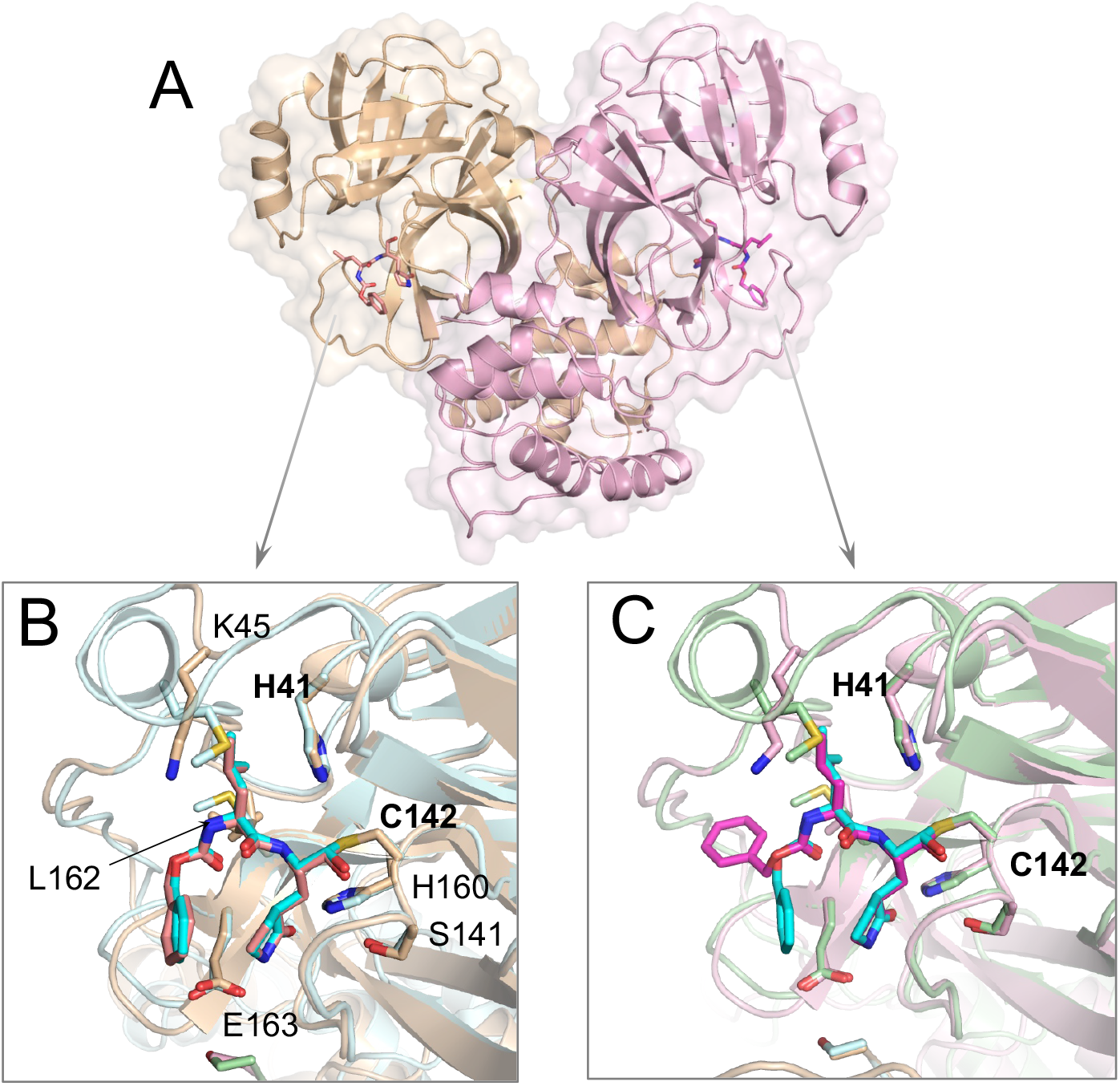
Crystal structure of M^pro^ from gamma-CoV SW1 bound to inhibitor GC376. (A) The overall structure is displayed above in cartoon representation with transparent surface and the two protomers colored pink and wheat. The inhibitor molecules covalently bound at the active sites are magenta and in stick representation. (B, C) The close-up views of the two active sites, where the SARS-CoV-2 M^pro^ structure (PDB 6WTJ) is superimposed and depicted in light blue and green cartoon representation. The key active sites residues interacting with the inhibitor are labeled in panel B. The terminal benzyl group of the inhibitor is flipped relative to the canonical binding mode (as seen in SARS-CoV-2 M^pro^; teal sticks) in one of the protomers (panel C).

The inhibitor GC376 was bound with the glutamine mimic g-lactam at the P1 position making direct interactions with Glu163, and inter-ligand stacking with the P3 benzyl ring in one of the protomers. At the active site of the other protomer, the P3 group was flipped such that the benzyl ring interacting with the protease instead of making inter-ligand interactions. A similar flipped binding mode of the P3 group for GC376 has been observed before in a crystal structure of this inhibitor with SARS-CoV-2 M^pro^ (PDB ID: 6WTT; [12]) although in most structures the benzyl ring stacks with the P1 group g-lactam (PDB IDs: 7JSU, 6WTJ; [11,13]). This dual binding mode of the P3 group suggests interactions at the S3 subsite could be optimized, ideally while maintaining inter-ligand interactions.

The leucine side chain at the P2 position was tucked in the hydrophobic S2 subsite, consistent with the substrate amino acid preference at this position. The P2 group interacts mostly with Glu186, located in the 180s loop. Amino acid sequence alignment of M^pro^ from various coronavirus species (Figure S1) shows that this position in the 180s loop is not conserved, with a Pro or Gln in alpha and beta-CoVs, respectively. Despite this variation, binding conformation of the P2 moiety was mostly the same in this gamma-CoV M^pro^ as in other reported structures, including SARS-CoV-2.

### Inhibitor binding to delta and gamma-coronavirus M^pro^

In addition to the gamma-CoV SW1, we determined the crystal structure of M^pro^ from the porcine delta-CoV HKU15 bound to PF-00835231. For both cocrystal structures, we quantified the contribution of active site residues to the packing with the inhibitor, by calculating the van der Waals interactions between the inhibitor and protease (Figure 2). These were compared with packing of the same inhibitor at the active site of SARS-CoV-2 M^pro^.

**Figure 2.**
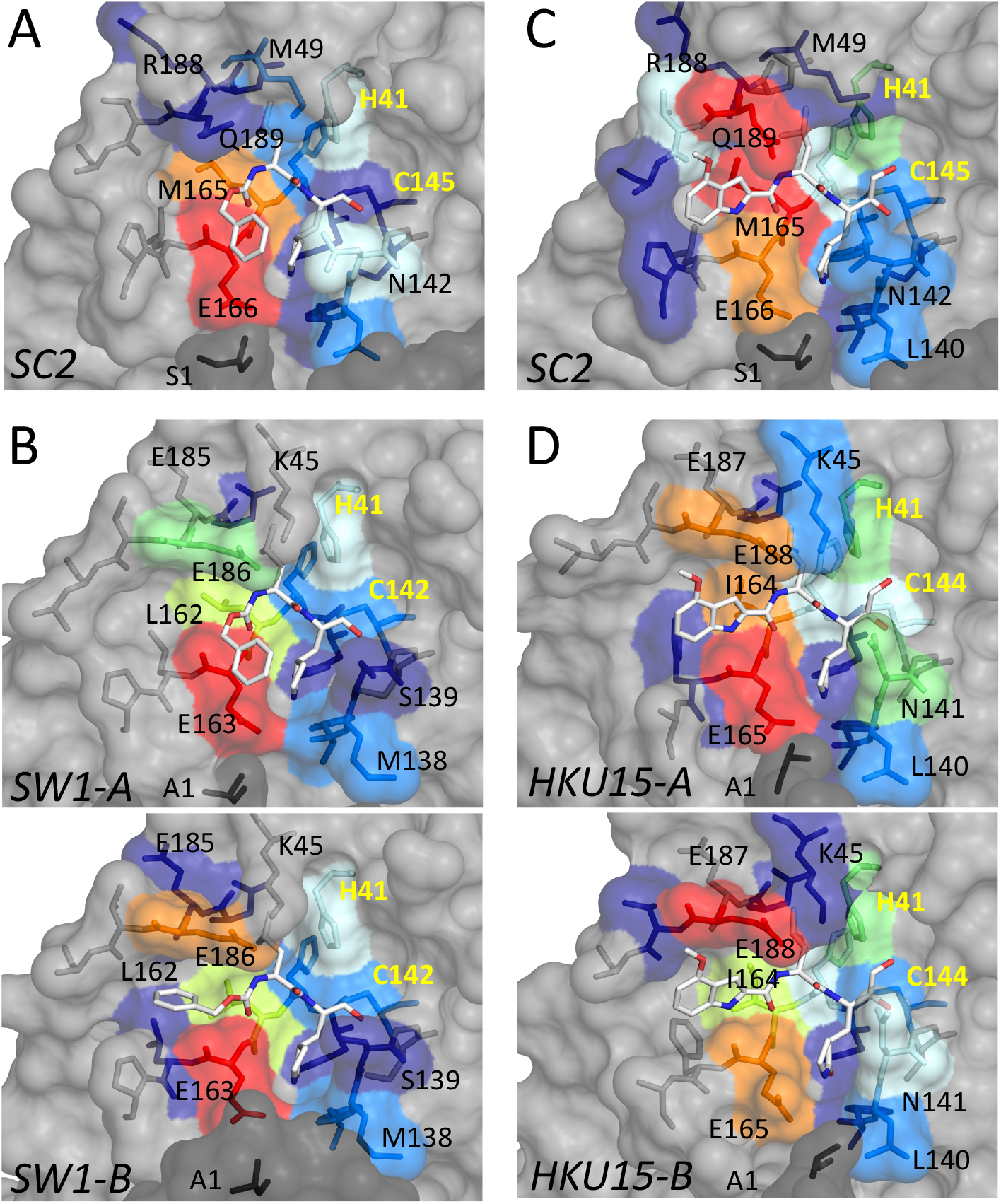
Packing of inhibitors with the active site residues in cocrystal structures of M^pro^ in comparison with SARS-CoV-2 (SC2). (A, B) Inhibitor GC376 bound to M^pro^ from SARS-CoV-2 (PDB ID: 6WTJ) and gamma-CoV SW1 from beluga whale. (C, D) Inhibitor PF-000835231 (active form PF-07304814) bound to M^pro^ from SARS-CoV-2 (PDB ID: 8DSU; presented in this work) and porcine delta-CoV HKU15. Interactions at the active sites of the two protomers are indicated by A and B for SW1 and HKU15. In all panels, the active site residues are colored blue to red for increasing interactions with the bound inhibitor. The catalytic residues are labeled in yellow font.

PF-00835231 bound at the active site of HKU15 M^pro^ with the same binding mode observed with SARS-CoV-2. Due to variation of amino acids at the active sites of two variants, the degree of packing with the inhibitor varied despite the same binding mode. Interactions with the 180s loop were slightly lower while those with the conserved Glu165/166 and N141/142 near the P1 group were higher in HKU15 M^pro^.

The P2 moiety of the inhibitors had extensive interactions with the Glu186/188. The leucine group of GC376 had considerably higher contacts with Glu186 of SW1 M^pro^, compared to Gln189 at the equivalent location in SARS-CoV-2 (Fig 2A,B). The 180s loop containing this Glu/Gln amino acid varies in conformation depending on the bound ligand (substrate peptide or inhibitor) in crystal structures reported [28]. Thus, the flexibility and variation in this loop might be relevant for accommodating different inhibitors.

Closer analysis of the determined cocrystal structures revealed that the E186/188 in the 180s loop establishes a salt bridge interaction with the side chain of K45 in both SW1 and HKU15 M^pro^ (Figure 3). Amino acid sequence alignment shows that these two amino acids are not conserved in alpha or beta-CoV (Figure S1), and the 40s loop is 3 amino acids shorter in delta- and gamma-CoV. Accordingly, there are no salt bridges in SARS-CoV-2 M^pro^ structures involving this relatively variable loop. Interestingly, this salt bridge was missing in the apo structure of SW1 M^pro^, suggesting that establishing of the salt bridge may correlate with ligand binding and/or stabilization of the 180s loop.

**Figure 3.**
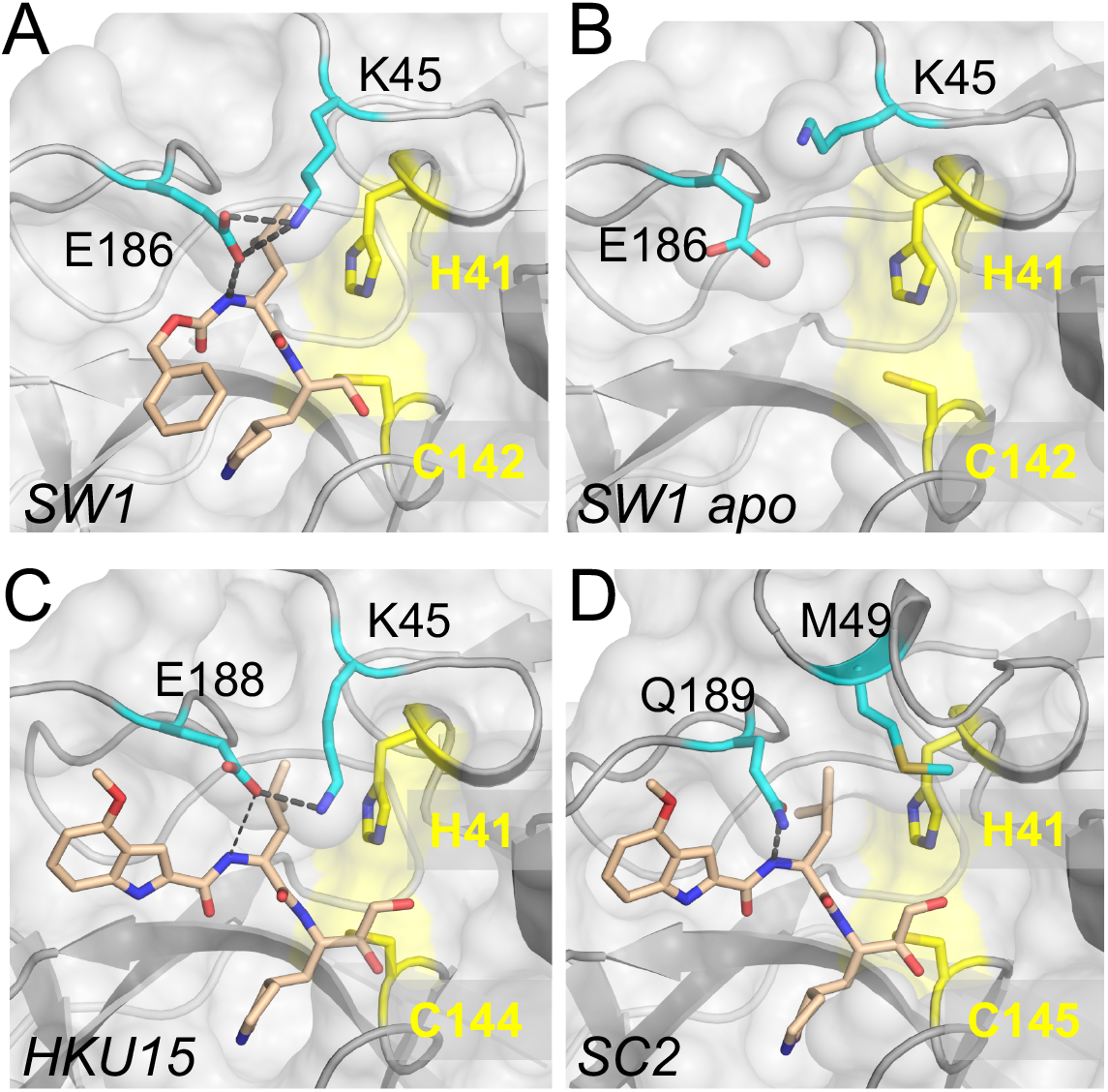
Salt bridge involving the 180s loop in delta- and gamma-CoV M^pro^ crystal structures. (A) Inhibitor-bound M^pro^ from beluga whale gamma-CoV SW1. The side chains of residues K45 and E186 establish a salt bridge over the P2 moiety of the bound inhibitor. (B) The salt bridge is missing in the apo structure of SW1 M^pro^. (C) Inhibitor PF-00835231 bound to M^pro^ from porcine delta-CoV HKU15. The salt bridge is between K45 and E188. (D) There are no residues that can form a similar salt bridge in alpha or beta-CoV M^pro^ variants, as shown for SARS-CoV-2 (SC2). In all panels, the protease is depicted in gray surface representation and the catalytic residues are colored yellow.

## Discussion

The abundance of coronavirus species in nature, especially in wild bats and birds, urge characterization of diverse coronavirus species that can adapt to infect domesticated animals and humans to cause outbreaks. Here we characterized the main protease of two highly divergent delta- and gamma-CoVs that have already adapted to infect mammals (Supplementary Figure 2). The M^pro^ from these two genera had been severely lacking in structural characterization; the crystal structures we present here constitute the first M^pro^ structures from a mammalian-infecting gamma-CoV. We found that the covalent small molecule inhibitors GC-376 and PF-00835231 bind to SW1 and HKU15 M^pro^ in their canonical binding mode, with relatively minor rearrangements, in agreement with their broad potency against various viral species.

The main differences among M^pro^ variants from the beta, delta, and gamma-CoV we analyzed here were in the S2 subsite where the amino acid sequences and structures are the most divergent (Figures 1-3 and S1). This subsite includes residues from the 180s loop, which is adaptable and changes conformation depending on the substrate peptide or inhibitor bound [28]. Residues in the 40s loop, which constitute the other part of the S2 subsite, are highly variable among CoV genera. These variations suggest the P2 position of inhibitors to be the most challenging to optimize to be able to effectively target all species. The shared leucine moiety at the P2 position of the inhibitors studied here is likely critical for their broad activity, and modifications at this position to increase potency against a given species may abrogate binding to other M^pro^ variants. Thus, our results suggest that the design of potential pan-coronavirus inhibitors need to maintain a leucine-like moiety at the P2 position. Our characterization of substrate peptide bound cocrystal structures of SARS-CoV-2 M^pro^ to determine the substrate envelope indicated that many of the current inhibitors protrude from the substrate envelope at this P2 position [28]. Such protrusions not only make these inhibitors susceptible to resistance mutations [28,30] but will also likely prevent achieving pan-coronaviral activity.

The M^pro^ from the gamma-CoV SW1 has the same overall fold as other 3C and 3C-like proteases, as expected, despite low sequence identity to the main proteases of alpha- and beta-CoV including SARS-CoV-2. Very few marine mammal-infecting gamma-CoVs have been identified to date: the whale-infecting SW1 whose M^pro^ we characterized here, and two species of dolphin-infecting coronaviruses [31,32]. Similarly, delta-CoVs are severely understudied. Both delta- and gamma-CoVs are thought to originate from birds and have already adapted to infect mammals [33]. The ability of coronaviruses to jump species barriers presents challenges for preventing and containing future outbreaks [33,34], signifying the need for pan-coronaviral treatment options. Considering the rapid evolution of the virus, especially of the spike protein to evade antibody and vaccine protection, M^pro^ is an attractive and relatively well-conserved target. Nevertheless, there is diversity among already known coronavirus species and characterization of their M^pro^ structures, as we have done here, should be useful in designing and developing effective pan-coronaviral inhibitors.

## Supplementary Materials

Figure S1: Amino acid sequence alignment of M^pro^ from various coronavirus species, Figure S2: Phylogenetic tree of coronaviruses, Table S1: Crystallization and refinement statistics.

## Data Availability

The data presented in this study are openly available in Protein Data Bank, accession codes 8DSU, 8FWX, 8E7N, 8E7C.

