## Supplementary Material for "Crystal Structures of Inhibitor-Bound Main Protease from Delta- and Gamma-Coronaviruses"

Table S1: Crystallization and refinement statistics of Mpro structures for SARS-CoV-2 (SC2), beluga whale (SW1) and swine (HKU15) coronaviruses.

| ***Mpro–Inhibitor*** | ***SC2–PF-00835231*** | ***SW1 apo*** | ***SW1–GC376*** | ***HKU15–PF-00835231*** |
|---|---|---|---|---|
| *PDB ID* | 8DSU | 8FWX | 8E7N | 8E7C |
| *Data Collection* | Home source | ALS 5.0.2 | Home source | Home source |
| *Resolution range (Å)*<br>*highest-res. shell* | 27.11-1.86<br>(1.926-1.86) | 70.59-2.57<br>(2.71-2.57) | 12.24-1.65<br>(1.709-1.65) | 22.91-2.45<br>(2.538-2.45) |
| *Space group* | P21 | P21212 | P21 | P61 |
| *a, b, c (Å)* | 54.9, 98.9, 59.1 | 82.3, 137.3, 49.8 | 63.1, 84.2, 68.5 | 64.1, 64.1, 261.8 |
| *α, β, γ (º)* | 90, 107.6, 90 | 90, 90, 90 | 90, 93.1, 90 | 90, 90, 120 |
| *Total reflections* | 242993 (15914) | 116131 | 199032 (15277) | 50645 (4137) |
| *Unique reflections* | 50477 (5011) | 18717 | 81531 (7698) | 20849 (1863) |
| *Multiplicity* | 4.8 (3.2) | 6.2(6.2) | 2.4 (2.0) | 2.4 (2.2) |
| *Completeness (%)* | 99.90 (99.60) | 99.9(99.9) | 94.95 (90.26) | 93.51 (83.92) |
| *Average I/σ* | 19.97 (1.85) | 11.3(2.3) | 11.31 (1.61) | 5.37 (1.26) |
| *Wilson B-factor* | 23.66 | 61.97 | 13.90 | 24.10 |
| $R_{merge}^{a}$ | 0.04739 (0.6403) | 0.126(0.966) | 0.05568 (0.4862) | 0.1608 (0.6731) |
| *CC1/2* | 0.999 (0.738) | 0.996(0.821) | 0.997 (0.735) | 0.973 (0.452) |
| *Refinement* | | | | |
| $R_{factor}^{c}$ | 0.2049 (0.3112) | 0.2297 | 0.1593 (0.2368) | 0.1927 (0.2385) |
| $R_{free}^{d}$ | 0.2543 (0.3744) | 0.2865 | 0.1951 (0.2747) | 0.2349 (0.2937) |
| *RMSD$^{b}$ in:* | | | | |
| *Bond lengths (Å)* | 0.004 | 0.009 | 0.018 | 0.002 |
| *Bond angles (º)* | 0.55 | 0.98 | 1.50 | 0.48 |
| *Ramachandran:* | | | | |
| *Favored (%)* | 97.35 | 97.16 | 98.48 | 94.19 |
| *Allowed (%)* | 2.65 | 2.84 | 1.52 | 5.13 |
| *Outliers (%)* | 0 | 0 | 0 | 0.68 |
| *Rotamer outliers (%)* | 0.99 | 0.85 | 0 | 3.85 |
| *B-factors:* | | | | |
| *Average* | 27.81 | 49.17 | 19.98 | 25.30 |
| *Macromolecules* | 26.54 | 49.05 | 17.56 | 24.98 |
| *Solvent* | 37.46 | 37.83 | 33.40 | 26.05 |

[a]Rsym = Σ | I − <I>|/ Σ I, where I = observed intensity, <I> = average intensity over symmetry equivalent.
[b]RMSD, root mean square deviation.
[c]Rfactor = Σ || Fo| − |Fc||/ Σ|Fo|.
[d]Rfree was calculated from 5% of reflections, chosen randomly, which were omitted from the refinement process.
Statistics for the highest-resolution shell are shown in parentheses.

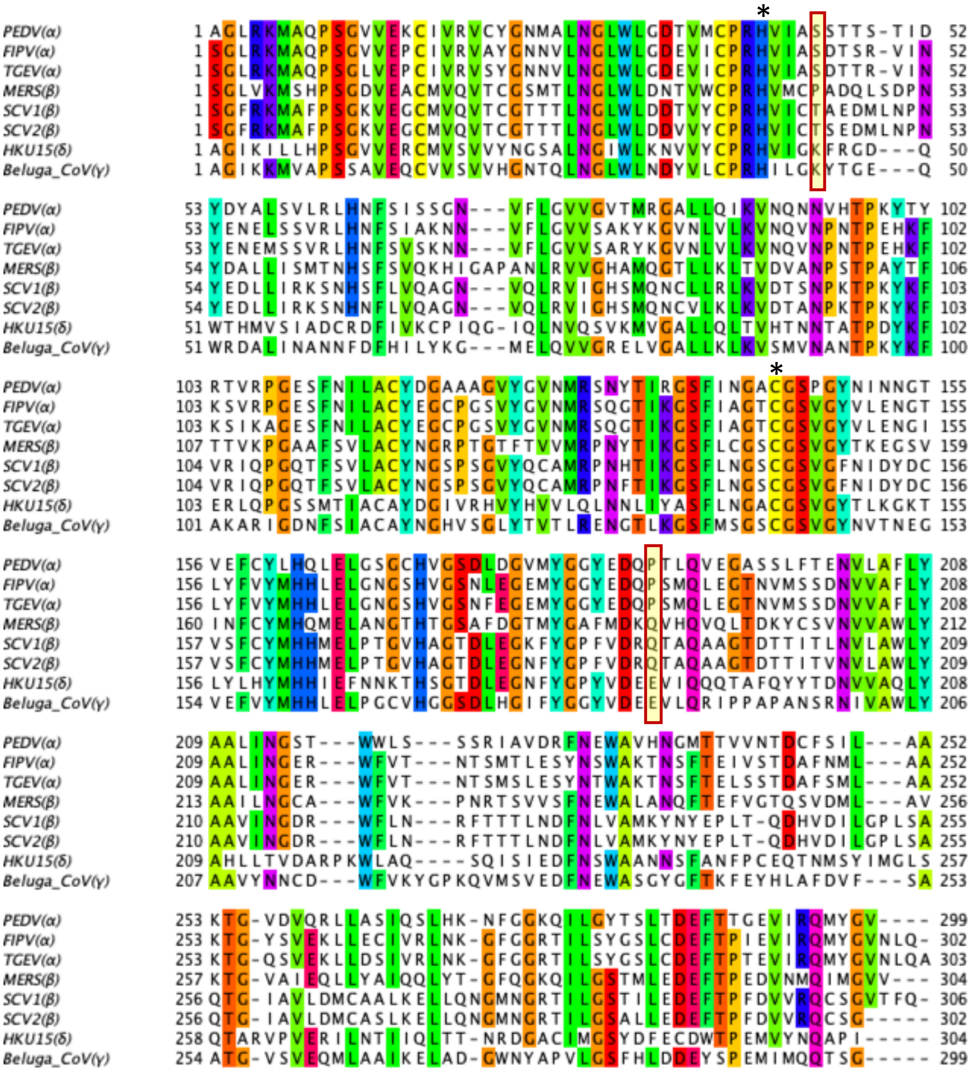

Figure S1. Amino acid sequence alignment of M$^{pro}$ from various coronavirus species. The conserved catalytic dyad residues (His and Cys) are indicated by asterisk, and the amino acids forming the salt bridge over the S2 subsite in delta- and gamma-CoV are highlighted in red frame. Alpha-CoVs are PEDV (porcine endemic diarrhea virus), FIPV (feline infectious peritonitis virus), TGEV (transmissible gastroenteritis virus); beta-CoV are MERS (Middle East respiratory syndrome), SCV1 (SARS-CoV-1), SCV2 (SARS-CoV-2), delta-CoV is the porcine HKU15 and the gamma-CoV is the beluga whale SW1. Multiple sequence alignment was performed using the Clustal web service with default parameters through the Jalview menu. The sequences are colored using the "Taylor" scheme with the threshold set at 60% identity.

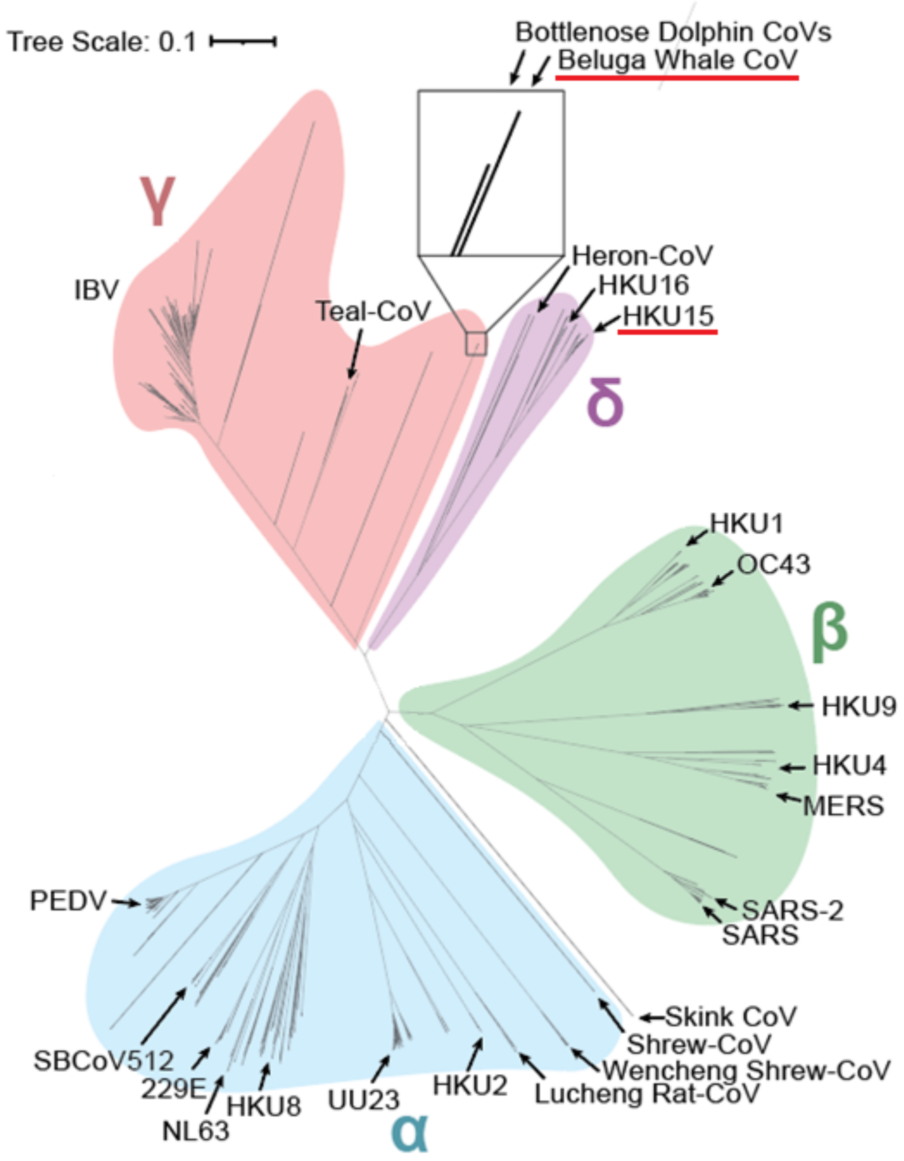


Figure S2. Phylogenetic tree of coronaviruses displaying the four main genera: alpha, beta, gamma, and delta. The tree was generated based on M$^{pro}$ amino acid sequences. The beluga whale (SW1) and porcine (HKU15) coronaviruses of which M$^{pro}$ structures were solved here are underlined.
